# Long-term Musculoskeletal and Marrow Sparing with Proton FLASH in Juvenile Mice: Implications for Pediatric Osteosarcoma

**DOI:** 10.64898/2026.01.16.699970

**Authors:** Zongsheng Hu, Uwe Titt, Yuting Li, Elise Konradsson, Edgardo A Aguilar, Denae W Neill, Luke Connell, Xiaochun Wang, Henry James Meyer, Mihai Lurascu Gagea, Susan L McGovern, Emil Schüler, Radhe Mohan

**Affiliations:** Department of Radiation Physics, University of Texas MD Anderson Cancer Center, Houston, Texas; Department of Veterinary Medicine & Surgery, University of Texas MD Anderson Cancer Center, Houston, Texas; Department of Radiation Oncology, University of Texas MD Anderson Cancer Center, Houston, Texas

**Author notes:** **Corresponding Author Name & Email Address:** Radhe Mohan.

## Abstract

**Purpose:** Osteosarcoma is the most common primary bone malignancy in children and adolescents. Radiotherapy is limited by intrinsic radioresistance and the risk of severe long-term musculoskeletal toxicities. FLASH radiotherapy, delivered at ultra-high dose rates (>40 Gy/s), has demonstrated normal tissue sparing in preclinical models, but its effects on the developing skeleton and marrow remain poorly defined. This study evaluated the chronic normal tissue effects of proton FLASH in juvenile mice, modeling the pediatric context.

**Methods and Materials:** Juvenile C57BL/6 mice (3–4 weeks old) were randomized to receive 11 Gy FLASH (≈200 Gy/s) or conventional proton irradiation (0.2 Gy/s average), or sham treatment to the left hind leg using a synchrotron-based proton beamline. Mice were followed for 10 weeks post-treatment. Bone toxicity was assessed with microCT (bone mineral density, bone volume fractions, trabecular indices) and histology. Bone marrow cellularity was quantified on H&E-stained sections, and muscle fibrosis was assessed using Masson’s trichrome.

**Results:** FLASH-treated mice exhibited significant preservation of bone microarchitecture compared with conventional treated mice, with higher bone mineral density and bone volume fractions (p < 0.05). Trabecular numbers were maintained, while structure model index indicates a mechanically favorable trabecular structures in the FLASH group. Bone marrow cellularity was preserved in FLASH mice (5.3% reduction vs. sham) compared with conventional (11.3 % reduction, p < 0.05). Muscle fibrosis was significantly lower in FLASH group (fibrosis positivity 2.6 % vs. 3.0% for FLASH vs. conventional CONV, p < 0.05). No severe immobility or weight loss was observed across groups.

**Conclusions:** Proton FLASH significantly reduces long-term bone, marrow, and muscle toxicities in juvenile mice. These findings provide the first demonstration of musculoskeletal sparing in a synchrotron proton FLASH platform and highlight its translational potential for pediatric osteosarcoma.

## Introduction

Osteosarcoma is the most common primary bone malignancy in children and adolescents, accounting for approximately 2.4% of cancers in this age group. The disease predominantly arises in the metaphysis of long bones, with the lungs and other bones as frequent sites of metastases. Prognosis is strongly determined by metastatic status: while patients with localized disease achieve 5-year survival rates of ∼77%, outcomes drop to ∼25% in those with distant metastases (1-4). Surgery remains the cornerstone of local control; however, it carries substantial morbidity and is often unfeasible for tumors in anatomically complex or axial locations such as the spine and pelvis (5-7). Radiotherapy offers a non-invasive alternative that preserves organ structure and function, yet the intrinsic radioresistance of osteosarcoma necessitates doses exceeding 64–70 Gy to achieve durable tumor control. Delivering such high doses with conventional radiotherapy is constrained by unacceptable toxicity to surrounding normal tissues, a limitation particularly pronounced in children, who are highly susceptible to growth impairment, deformities, and late tissue complications (8-12). Thus, innovative approaches that optimize therapeutic efficacy while minimizing collateral toxicity to adjacent normal tissues are of vital importance.

FLASH radiotherapy, delivered at ultra-high dose rates (>40 Gy/s), has emerged as a promising modality capable of reducing normal tissue toxicities without compromising tumor control. Extensive preclinical work across multiple organs and irradiation modalities—including lung, brain, intestine, and skin—has consistently demonstrated the potential of FLASH to widen the therapeutic window (13-23). Both electron and proton beams have been shown to produce the FLASH normal tissue sparing effect; however, electrons are restricted by their shallow penetration, limiting application to superficial targets. Protons, in contrast, combine clinically relevant depth penetration with the dosimetric advantages of the Bragg peak, enabling conformal irradiation while minimizing dose to adjacent normal tissues. These physical properties position proton FLASH as a promising candidate for clinical translation to pediatric osteosarcoma.

A major concern for radiotherapy in osteosarcoma is chronic toxicity to bone and muscle, as these tissues are essential for growth, mobility, and long-term function in pediatric patients. Preclinical studies using murine models have provided encouraging evidence that proton FLASH has the potential to mitigate such toxicities. Proton pencil beam scanning FLASH has been shown to reduce leg contracture and skin damage (24), while additional studies demonstrated preservation of bone and muscle integrity with maintained sarcoma control (25). More recently, proton FLASH reirradiation further alleviated late effects including fibrosis, lymphedema, and tibial fractures (26). These findings suggest that proton FLASH may offer meaningful protection to the very tissues most vulnerable in osteosarcoma treatment.

However, critical gaps remain. Nearly all prior investigations have focused on adult mice models (∼6–13 weeks), neglecting the unique vulnerabilities of the developing skeleton and marrow compartment in pediatric patients. In addition, long-term effects on bone integrity, marrow reserve, and muscle for juvenile models remain insufficiently characterized, despite their relevance as dose-limiting tissues in pediatric patients. Furthermore, most all existing proton FLASH studies have been performed with cyclotron-based systems, leaving it uncertain whether the FLASH effect can be reproduced with synchrotron platforms, which are also widely used in hospital-based proton therapy facilities worldwide.

In this study, we address these biological gaps by evaluating the chronic effects of synchrotron-based proton FLASH irradiation in juvenile mice (3–4 weeks old), a developmental stage that mirrors the pediatric patient context. Using micro-CT and histopathological analyses, we demonstrate that FLASH irradiation significantly reduces long-term toxicities in bone, bone marrow, and muscle compared with conventional dose rates. To our knowledge, this represents the first demonstration of sustained musculoskeletal sparing in a synchrotron-based proton FLASH platform. These findings establish biological evidence of supporting proton FLASH as a compelling strategy to reduce morbidity and improve outcomes in pediatric osteosarcoma. If translatable to patients, such sparing could promote preservation of skeletal development, reduce long-term morbidity, and expand the role of radiotherapy as a definitive treatment option for unresectable pediatric osteosarcoma.

## Methods

### Animal Model and Experimental Design

All procedures were conducted in accordance with institutional guidelines and approved by the Institutional Animal Care and Use Committee (IACUC). Healthy female juvenile C57BL/6 mice aged 3–4 weeks (weight range: (6 – 11 g) were selected to model the pediatric context of osteosarcoma patients and randomly assigned to treatment groups. Animals were housed in specific pathogen–free facilities with controlled temperature, humidity, and a 12-hour light/dark cycle, with ad libitum access to food and water.

Mice were divided into three groups: FLASH irradiation (FLASH; n = 14), conventional irradiation (CONV; n = 12), and sham-irradiated controls (SHAM; n = 9). Each mouse received unilateral irradiation to the left hind leg, with the contralateral leg serving as an internal control. Body weight and clinical condition (posture, grooming, activity) were recorded weekly. Protocol-defined endpoints included >20% body weight loss, impaired mobility, or ulceration requiring euthanasia. All mice were followed longitudinally for 10 weeks post-irradiation to assess chronic toxicities. At the study endpoint, mice were euthanized by CO₂ inhalation followed by cervical dislocation, and tissues were harvested for analysis.

### Proton Irradiation and Dosimetry

#### Anesthesia, Positioning, and Irradiation Setup

Mice were anesthetized with 2% isoflurane for induction in the chamber and 1.5% for maintenance in the nose cone with ambient air as carrier gas and secured in a custom-designed immobilization jig to ensure reproducible positioning of the hind limb (Figure 1c). A 3D-printed helmet stabilized the head and provided continuous delivery of isoflurane during irradiation. The left leg was gently extended and taped in place so that the femur and tibia were fully encompassed within the treatment field. A lead shield contoured to the aperture minimized off-target exposure; the circular high-dose field is indicated by the red circle in Figure 1c, corresponding to the region receiving ≥90% of the prescribed dose.

**Figure 1.**
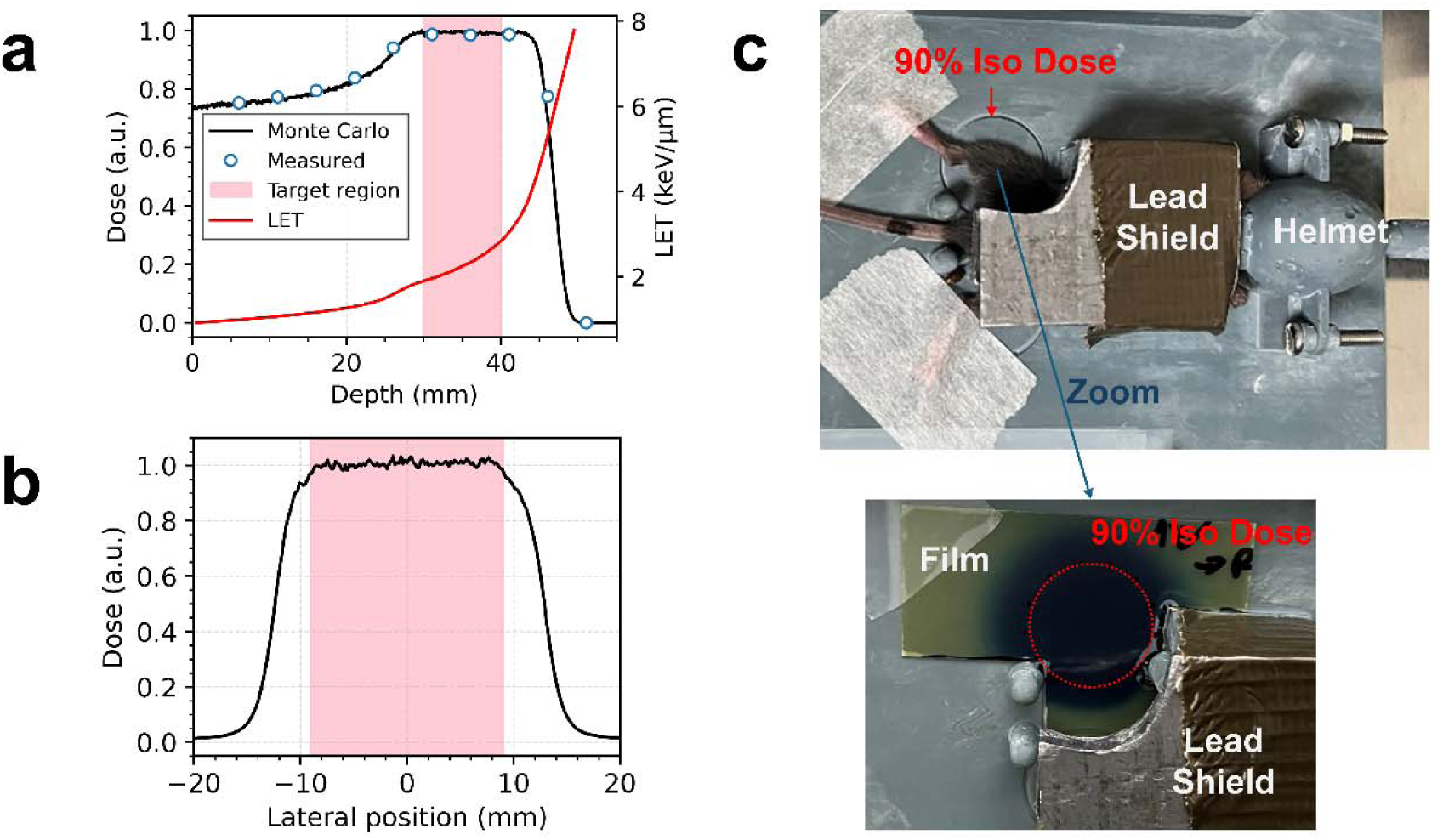
(a) Depth-dose distribution for the 2-cm SOBP: ion chamber measurements (symbols) compared with Monte Carlo simulations (solid line), with the corresponding linear energy transfer (LET) curve shown in red. (b) Lateral dose profile across the 2 cm diameter irradiation field obtained with EBT3 Gafchromic film, confirming field flatness. (c) Photograph of the custom mouse immobilization jig with the 3D-printed helmet for isoflurane delivery. The red circle marks the region receiving ≥90% of the prescription dose; a lead shield restricts exposure outside the target field.

Irradiations were performed at a synchrotron-based proton beamline manufactured by Hitachi. A 2-cm spread-out Bragg peak (SOBP, see Figure 1a and 1b) was created and delivered through a 2-cm-diameter circular field to the left hind limb. A 3-cm water-equivalent pre-absorber was positioned upstream to ensure that the target volume was centered within the SOBP with a dose averaged LET of 2.1 keV/µm. The prescribed dose of 11 Gy was delivered in a single fraction. Absolute dosimetry was carried out using an Advanced Markus Chamber. The recombination factor was determined using the standard 2-voltage approach (+300 and +150 V) recommended in the TRS-398 report (27). Beam output was monitored online with transmission chamber cross-calibrated against the reference chamber. Beam uniformity across the 2 cm- diameter field at 3.5 cm depth was confirmed before each irradiation with EBT3 Gafchromic films (Ashland, USA) (Figure 1b). Depth-dose distributions were measured with the reference chamber and compared to Monte Carlo simulations for verification of SOBP range and flatness (Figure 1a). The beam’s temporal structure was validated using a digital oscilloscope connected to the beam current monitoring system, ensuring accurate reproduction of both FLASH and CONV delivery modes. All dosimetric checks confirmed delivery accuracy within ±5%.

FLASH delivery was achieved with a single synchrotron spill, resulting in an average dose rate of ∼200 Gy/s at isocenter (28). CONV irradiation was delivered using 26 consecutive spills, each delivering 0.64 Gy at an instantaneous rate of 1.28 Gy/s, separated by ∼1.5 s intervals, achieving a mean dose rate of ∼0.2 Gy/s. Sham animals underwent identical anesthesia and positioning procedures but received no irradiation.

### MicroCT

Imaging was performed 10 weeks post-irradiation using a Bruker micro-CT SkyScan 1276 (Bruker, Kontich, Belgium) at an isotropic voxel size of 13 μm, with tube voltage 55 kV and current 200 μA. Backward projection datasets of all the mice were reconstructed using Insta-Recon software (Bruker microCT, Kontich, Belgium). For the femur, the trabecular region of interest (ROI) was defined beginning 50 slices (∼650 μm) proximal to the growth plate and extending across 150 slices (∼1.95 mm); for the tibia, the ROI was defined as 50 slices distal to the growth plate and extended over 100 slices (∼1.3 mm). The analysis workflow is summarized in Figure 2a. First, the trabecular compartment was segmented from cortical bone, and histogram analysis of this region was performed to derive bone mineral density (BMD). Next, trabecular bone structures were segmented from the trabecular region using a consistent gray-value threshold across all samples to ensure objectivity. Each segmentation was manually reviewed to confirm accuracy. Representative examples of segmented trabecular bone in femur from CONV-irradiated, FLASH-irradiated, and sham-irradiated (SHAM) mice are shown in Figure 2b. The resulting binary masks of trabecular bone were used for morphometric analysis, including bone volume fraction (BV/TV), trabecular number (Tb.N), connectivity density (Conn.D), and structure model index (SMI). All quantitative parameters were calculated using the CTan software (Bruker, Kontich, Belgium).

**Figure 2.**
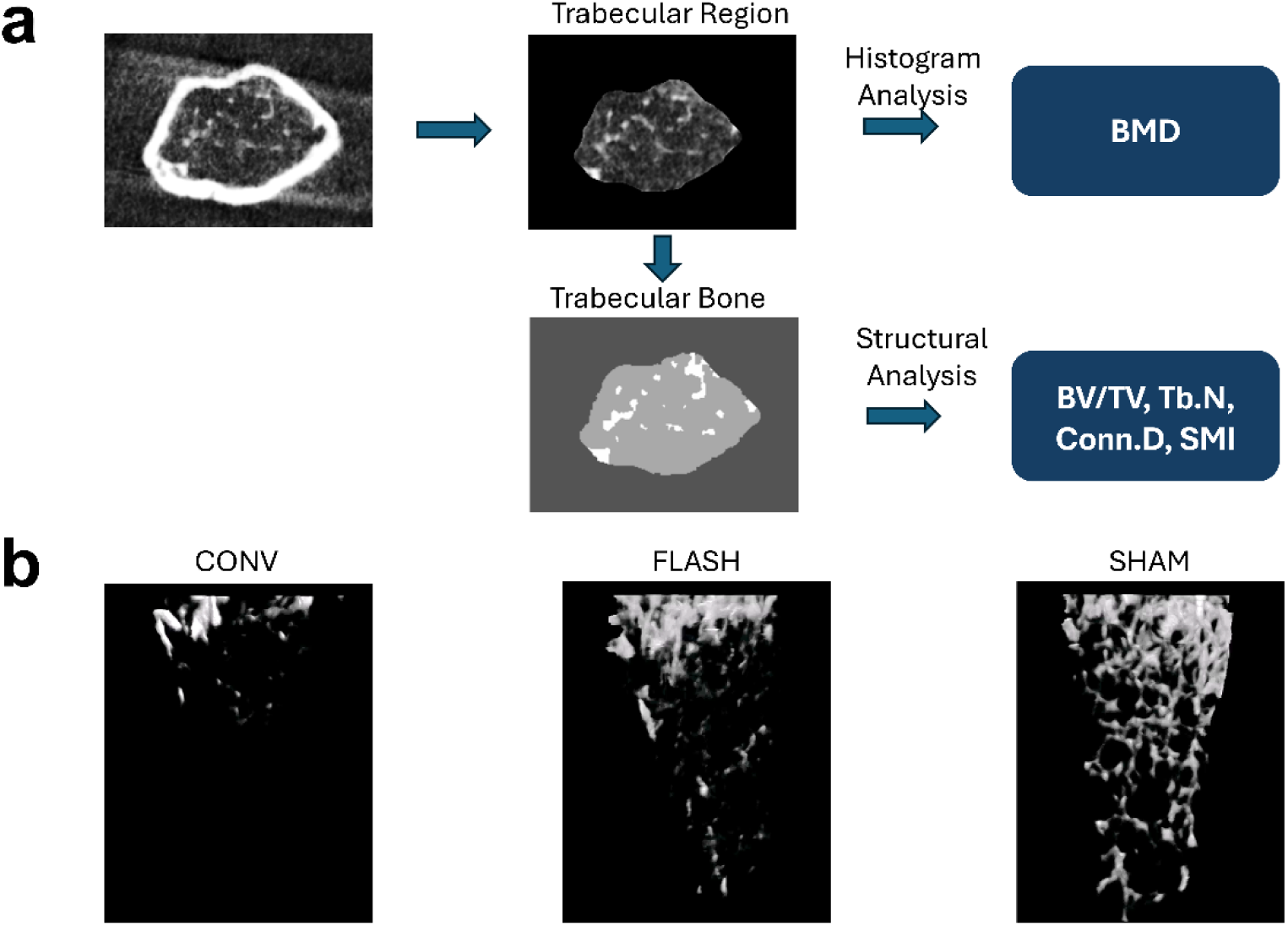
(a) Workflow of trabecular bone analysis. The trabecular compartment was segmented from cortical bone for histogram analysis (yielding bone mineral density, BMD). Trabecular bone was then filtered from the trabecular region and used for structural analysis (bone volume fraction, BV/TV; trabecular number, Tb.N; Connectivity Density, Conn.D; and structure model index, SMI). (b) Representative examples of segmented trabecular bone structure from conventionally irradiated (CONV), FLASH-irradiated, and sham-irradiated (SHAM) mice at 10 weeks post-treatment.

### Histopathology

At 10 weeks post-irradiation, femora, tibiae, and gastrocnemius muscles were harvested for histological analysis. Bones were decalcified in 10% ethylenediaminetetraacetic acid (EDTA), paraffin-embedded, and sectioned longitudinally for hematoxylin and eosin (H&E) staining to evaluate bone marrow cellularity. Stained sections were digitally scanned using a Versa 8 slide scanner at 20× magnification. The bone marrow region was manually segmented, and irrelevant areas (e.g., cracks or artifact bubbles) were excluded. Bone marrow cellularity was quantified as the number of nucleated hematopoietic cells per mm² of marrow space using the nuclear count algorithm in Aperio ImageScope (Leica Biosystems).

Skeletal muscle samples were fixed in 10% neutral-buffered formalin, embedded in paraffin, and sectioned both cross-sectionally and longitudinally at 5 µm thickness. Sections were stained with Masson’s trichrome to visualize radiation-induced fibrosis and scanned using the same imaging protocol and magnification. Collagen deposition was quantified as the percentage of collagen-positive area relative to total muscle area using the Positive Pixel Count algorithm in Aperio ImageScope, with non-muscle structures such as fat, vessels, and nerves excluded by manual segmentation.

Standard histological methods were applied across all tissue specimens: H&E staining was used for structural evaluation and cell density assessment, and trichrome staining was used to assess fibrosis. All image analyses were performed with investigators blinded to treatment groups.

### Statistical Analysis

Data are expressed as mean ± standard deviation unless otherwise stated. Normality was tested using the Shapiro–Wilk test. For normally distributed data, comparisons were made with unpaired two-tailed Student’s T-tests or one-way ANOVA with Tukey’s post hoc test. For non-normally distributed data, Mann–Whitney U tests were applied. A p-value <0.05 was considered statistically significant. Statistical analyses were performed with Python using SciPy package. Sample size was based on prior FLASH studies (24–26), which demonstrated significant tissue sparing with group sizes of 5–15 animals. Blinding was maintained during histological analysis and data quantification.

## Results

### General Observations and Animal Tolerability

All mice tolerated the irradiation procedures without acute morbidity. No procedure-related deaths occurred during the 10-week follow-up. Body weight remained stable across groups, with no significant differences between FLASH, CONV, and SHAM cohorts throughout the study period (Figure 5c). Health monitoring, including daily inspection for grooming, posture, and activity, revealed no overt signs of distress. Limb mobility was preserved in all groups, and no severe dermatitis, ulceration, or immobility was observed. These findings confirm that both FLASH and CONV irradiation were tolerated at the delivered dose of 11 Gy.

### FLASH Preserves Bone Microarchitecture

Figure 2b shows representative segmented trabecular structures from the three treatment groups. By visual inspection, bones from the FLASH group displayed a denser and more interconnected trabecular network compared with the CONV group, suggesting preservation of structural integrity. Sham animals demonstrated the most robust architecture, as expected. Quantitative microCT results are summarized in Figures 3 and 4 for the femur and tibia, respectively. In both sites, FLASH-treated mice consistently outperformed CONV irradiation in measures of trabecular preservation.

**Figure 3.**
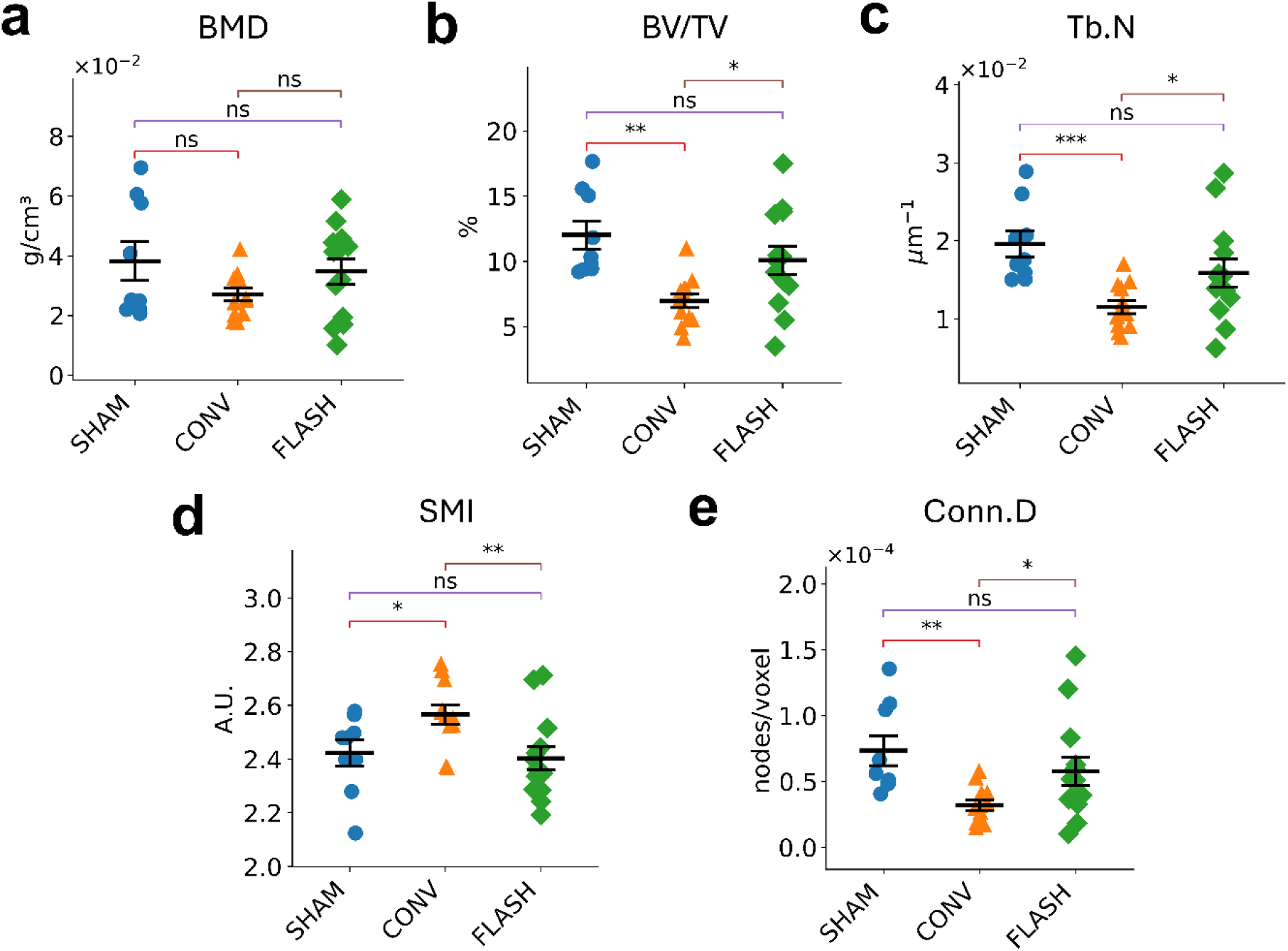
Quantitative microCT analysis of femoral trabecular bone at 10 weeks post-irradiation. FLASH-treated femora demonstrated significantly higher, bone volume fraction (BV/TV, b), trabecular number (Tb.N, c), and connectivity density (Conn.D, e) compared with conventional irradiation (CONV). Structure model index (SMI, d) was significantly lower in the FLASH group, consistent with a more plate-like morphology. Sham animals exhibited the best overall values across parameters (*p < 0.05, **p < 0.01, ***p < 0.001, ns p>0.05).

**Figure 4.**
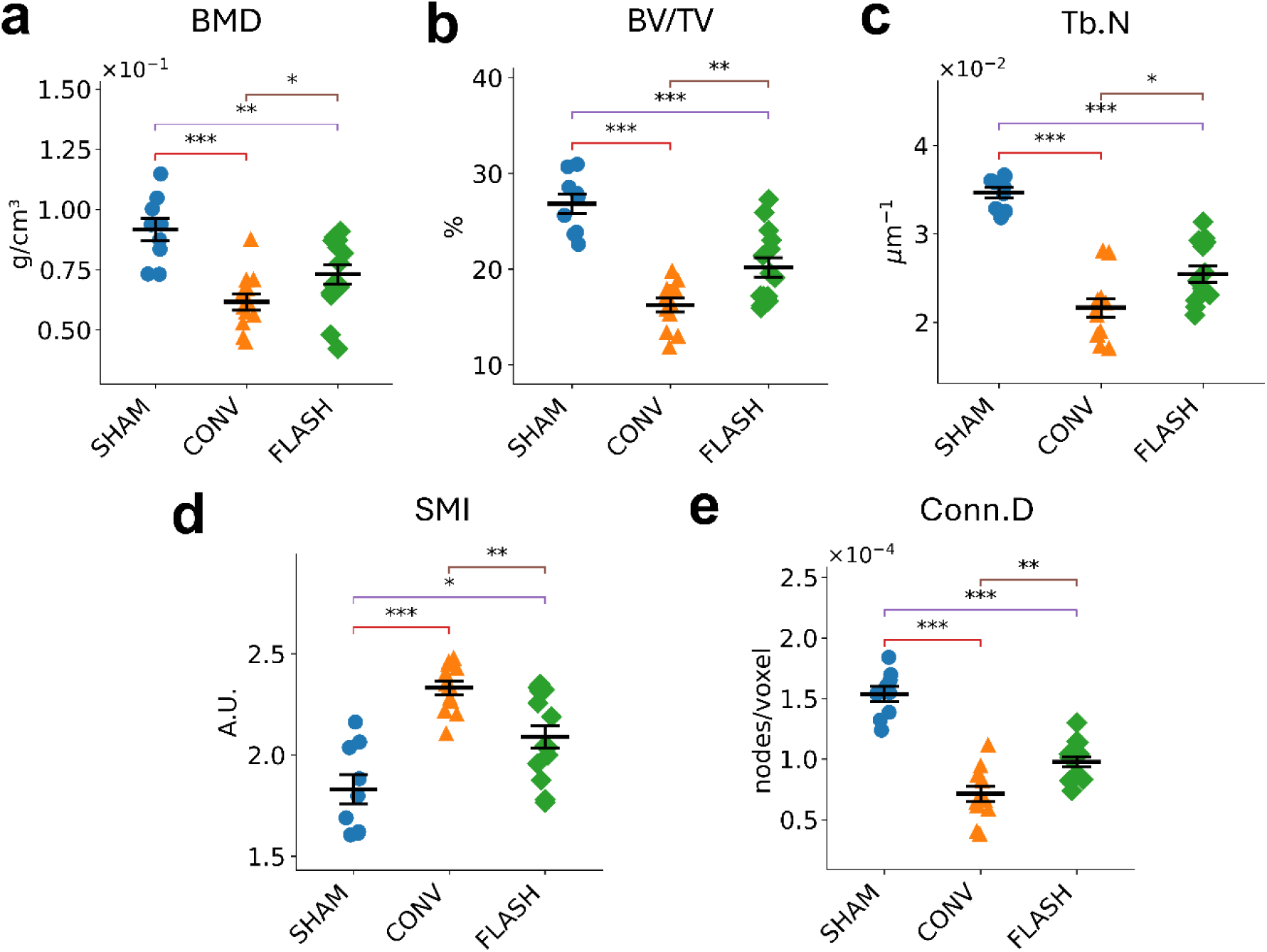
Quantitative microCT analysis of tibial trabecular bone at 10 weeks post-irradiation. Results paralleled those in the femur, with FLASH-treated tibiae showing significantly higher BMD (a), BV/TV (b), and Tb.N (c), as well as significantly lower SMI (d) and greater Conn.D (e) compared with conventional irradiation (CONV). Sham animals again exhibited the best overall bone structure (*p < 0.05, **p < 0.01, ***p < 0.001, ns p>0.05).

For bone mineral density (BMD), FLASH-treated tibiae maintained significantly higher values than CONV irradiated bones, approaching those of sham controls. Bone volume fraction (BV/TV) was likewise significantly greater for both femur and tibiae following FLASH, indicating preservation of trabecular structure within the trabecular compartment. Trabecular number (Tb.N), which quantifies the number of trabeculae per unit volume, was elevated in the FLASH group compared with CONV irradiation. Structure model index (SMI) values were significantly lower in FLASH-treated samples, consistent with preservation of a plate-like morphology, indicating a stronger and mechanically favorable shape, whereas CONV irradiation produced a shift toward a rod-like architecture associated with reduced mechanical strength(29). Finally, connectivity density (Conn.D), which quantifies the number of connections (nodes) between trabeculae per unit volume, was significantly improved in the FLASH group relative to CONV, indicating maintenance of trabecular interconnections. Together, these findings confirm that FLASH irradiation mitigated radiation-induced deterioration of trabecular bones and preserved microarchitectural complexity compared to CONV radiotherapy.

### FLASH Maintains Bone Marrow Cellularity

Histological analysis of H&E-stained sections revealed differences in bone marrow preservation between treatment groups (Figure 5a). In CONV irradiated mice, marrow cavities appeared hypocellular, with marked depletion of hematopoietic elements and frequent replacement by adipocytes, consistent with chronic radiation-induced marrow toxicity. In contrast, FLASH-treated animals maintained substantially higher cellular density, with marrow spaces populated by numerous nucleated precursors.

**Figure 5.**
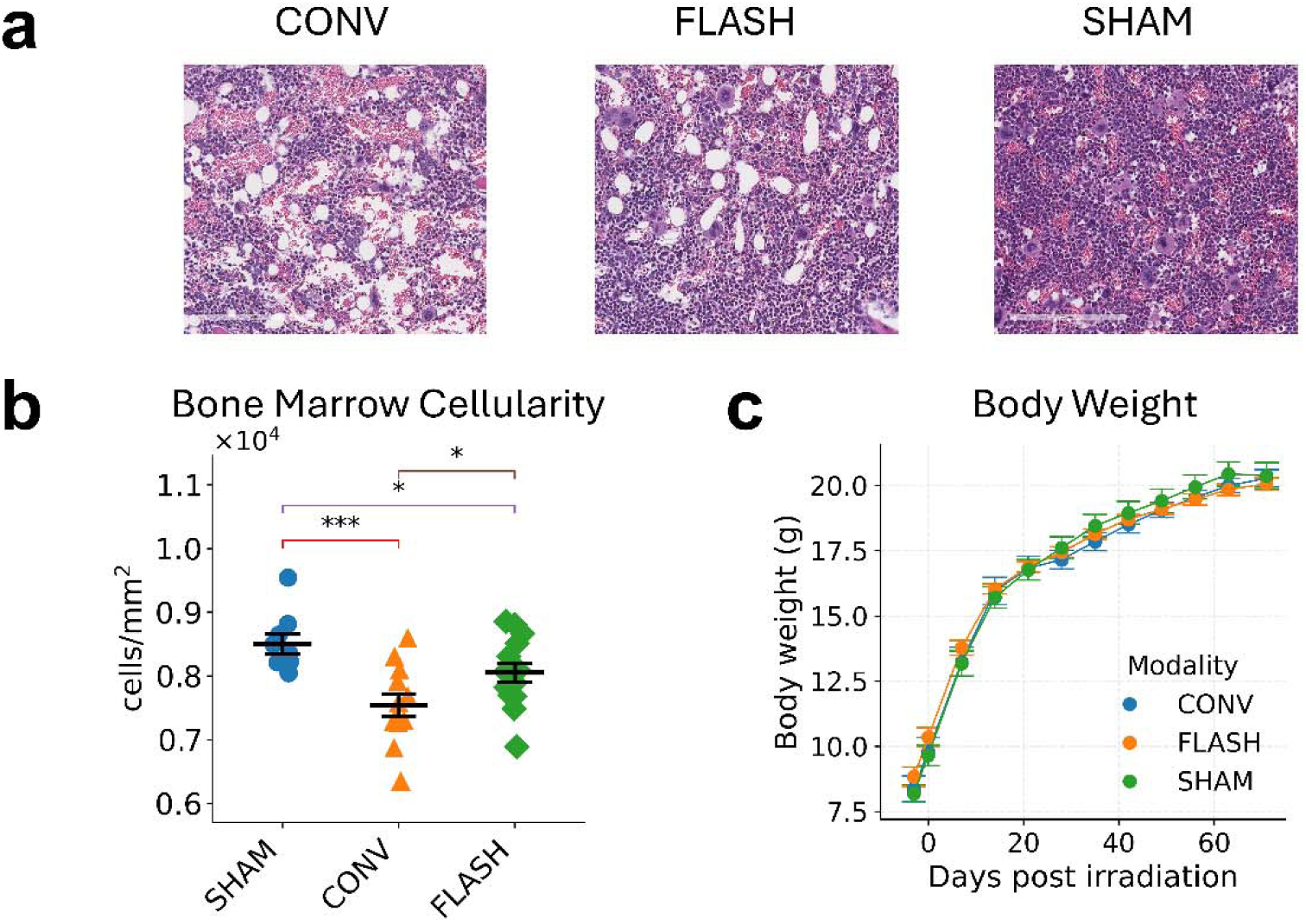
(a) Representative H&E-stained bone marrow sections from conventionally irradiated (CONV), FLASH-irradiated, and sham-irradiated (SHAM) mice at 10 weeks post-treatment. Conventional irradiation produced hypocellular marrow with adipocytic replacement, whereas FLASH preserved dense hematopoietic precursors more similar to sham. (b) Quantitative analysis of marrow cellularity (cells/mm²). Conventional irradiation significantly reduced marrow cellularity compared with sham, while FLASH animals showed significantly less reduction (*p < 0.05, **p < 0.01, ***p < 0.001, ns p>0.05). (c) Body weight curves of mice in each group during the 10-week follow-up period.

Quantitative image analysis confirmed these observations (Figure 5b). Compared with sham animals, CONV irradiation led to an approximate 11.3% reduction in marrow cellularity (p < 0.001), while FLASH-treated animals exhibited only about 5.3% reduction, a difference that reached statistical significance compared with the CONV group (p < 0.05). These findings were consistent across both femoral and tibial compartments. The preservation of marrow cellularity in the FLASH cohort suggests that ultra-high dose-rate irradiation spares critical hematopoietic progenitor populations that are otherwise highly sensitive to radiation damage, which remains in the long term.

### FLASH Reduces Muscle Fibrosis

Masson’s trichrome staining revealed clear differences in radiation-induced fibrosis among the treatment groups (Figure 6a). In CONV irradiated mice, both cross-sectional (top panels) and longitudinal (bottom panels) images of gastrocnemius muscle showed abundant collagen deposition (blue staining), with thickened interstitial septa and disorganized myofiber architecture, consistent with radiation-induced fibrosis. By contrast, FLASH-treated animals exhibited markedly reduced collagen accumulation, with preservation of normal myofiber structure and thinner, well-aligned collagen bundles. Sham-irradiated controls displayed uniformly intact fibers with minimal collagen deposition.

**Figure 6.**
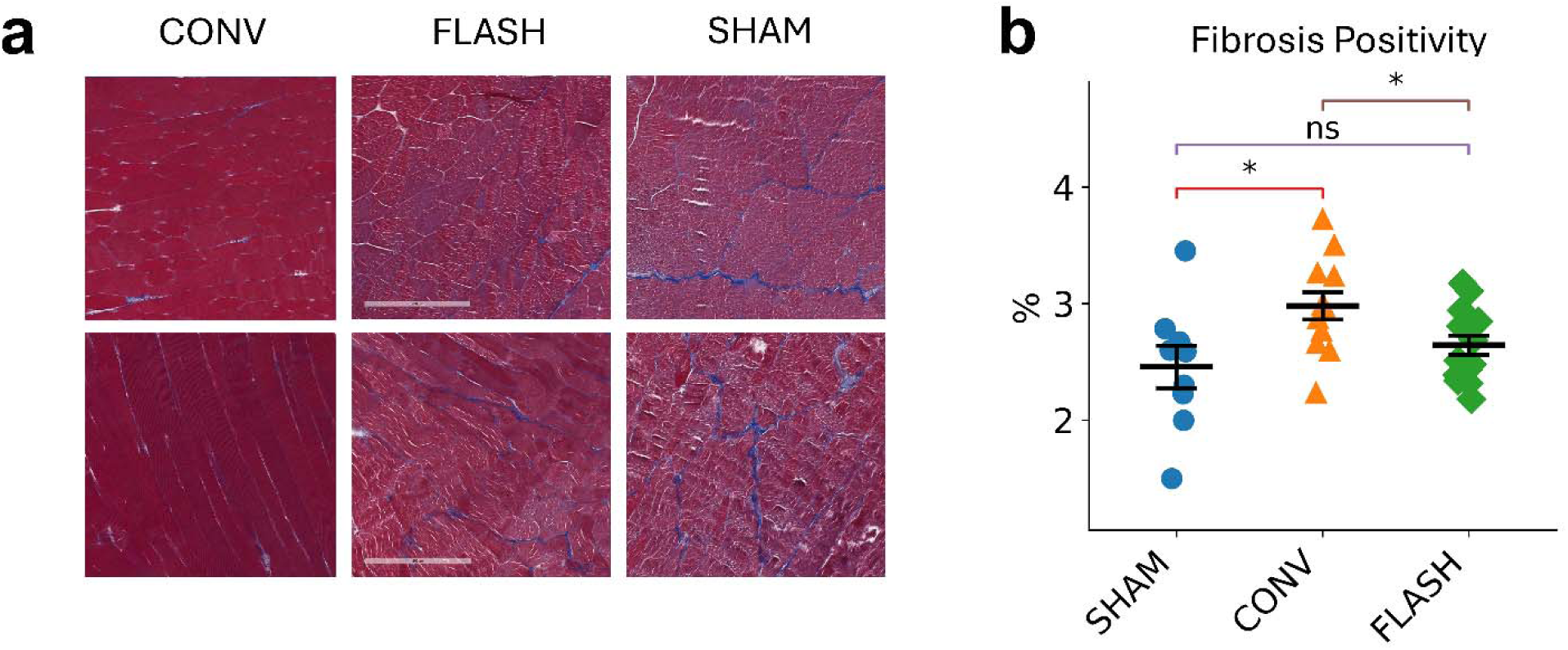
(a) Representative Masson’s trichrome-stained sections of gastrocnemius muscle at 10 weeks post-irradiation. Top panels show cross-sections, and bottom panels show longitudinal sections. Conventional irradiation (CONV) produced marked collagen deposition (blue staining) with thickened collagen between myofibers, whereas FLASH-treated muscles displayed reduced collagen accumulation and preserved fiber organization. Sham muscles showed minimal collagen deposition and intact morphology. (b) Quantification of fibrosis positivity (% collagen-positive area/total muscle area). FLASH significantly reduced fibrosis compared with conventional irradiation, with values closer to sham (*p < 0.05, **p < 0.01, ***p < 0.001, ns p>0.05).

Quantitative image analysis confirmed these qualitative impressions (Figure 6b). Fibrosis positivity was significantly lower in FLASH-irradiated muscles compared with CONV irradiation (3% vs. 2.6%, p < 0.05), and values in the FLASH group are closer to those observed in sham controls. These results demonstrate that FLASH not only reduces collagen deposition but also maintains muscle architecture, mitigating chronic radiation-induced fibrosis.

## Discussion

In this study, we demonstrate that synchrotron-based proton FLASH irradiation significantly reduces chronic toxicities in bone, bone marrow, and muscle in juvenile mice compared with CONV dose-rate irradiation. Using microCT and histopathological analyses, we observed preservation of bone mineral density, trabecular structure, marrow cellularity, and muscle architecture following FLASH delivery. To our knowledge, this represents the first report of sustained musculoskeletal sparing achieved with a synchrotron FLASH platform. These findings directly address critical concerns in pediatric osteosarcoma, where late toxicities in developing bone and muscle limit the therapeutic window of radiotherapy.

Our findings demonstrate that proton FLASH irradiation confers durable protection against late musculoskeletal toxicities in juvenile mice. Compared with CONV irradiation, FLASH preserved trabecular bone microarchitecture, maintained bone marrow cellularity, and mitigated muscle fibrosis. These results establish both the biological relevance and the technical feasibility of synchrotron-delivered proton FLASH, underscoring its translational potential as a strategy to improve therapeutic outcomes and reduce long-term morbidity in pediatric osteosarcoma.

Our results align with and extend previous evidence of normal tissue sparing with proton FLASH. Cunningham and colleagues reported that proton pencil beam scanning FLASH reduced leg contracture and skin toxicity in mice (24). Velalopoulou et al. further demonstrated protection of muscle and bone integrity while maintaining sarcoma control, implicating FLASH as a potential modality for mesenchymal tumors (25). More recently, Verginadis et al. showed that proton FLASH reirradiation of the mouse leg alleviated late effects including fibrosis, lymphedema, and tibial fractures, highlighting its capacity to mitigate chronic musculoskeletal toxicities under cumulative dose conditions (26). Importantly, all of these studies were performed in adult animals (6–13 weeks old) using cyclotron-based systems. By contrast, we evaluated juvenile mice (3–4 weeks old), modeling the pediatric clinical context, and employed a synchrotron delivery platform. Thus, our findings both confirm the reproducibility of the FLASH effect across different delivery technologies and uniquely establish its relevance to the developing skeleton.

Interestingly, our results differ from those of Liu et al. (30) who investigated acute gastrointestinal toxicity using the same synchrotron-based proton beamline. In their study, proton FLASH unexpectedly exacerbated intestinal toxicity compared with CONV dose rates, whereas linac-based electron FLASH conferred the expected sparing effect. Some other studies using various proton FLASH beamlines also show lack of sparing or even increased GI toxicity under abdominal FLASH conditions (31,32). In contrast, we observed a clear protective effect of proton FLASH in bone, bone marrow, and muscle, where late toxicities were significantly reduced relative to CONV irradiation. These contrasting outcomes underscore that the FLASH effect may not be uniform across tissues and that organ-specific biology likely determines the dose and dose rate threshold for benefit versus harm. The gastrointestinal tract, with its rapid epithelial turnover, high oxygenation, and dependence on stem cell niche regeneration, may require different dose and dose rate conditions to achieve FLASH effect than musculoskeletal tissues, which are characterized by slower turnover, greater hypoxic fractions, and fibrotic injury pathways. Together, these studies underscore that the dose and dose-rate thresholds required to reach the FLASH effect are tissue dependent and that proton FLASH may be beneficial in some organs (e.g., bone and muscle) but potentially harmful in others (e.g., intestine) under the same beam conditions. Mechanistic studies dissecting the interplay between tissue oxygenation, repair kinetics, and radiation response will be critical to guiding safe and effective clinical application of FLASH therapy.

Muscle and bone are generally regarded as late-responding tissues, in which radiation injury evolves gradually over weeks to months. Muscle toxicity is dominated by progressive fibrotic remodeling and myofiber atrophy, while bone damage is characterized by long-term deterioration of trabecular structure, cortical thinning, and increased fracture risk. In our study, proton FLASH significantly reduced both muscle fibrosis and trabecular bone degradation at 10 weeks, indicating that ultra-high dose-rate delivery can mitigate these late toxicities. Clinically, such protection is highly relevant in pediatric osteosarcoma, where late musculoskeletal complications can compromise growth, mobility, and quality of life long after initial treatment. The ability of FLASH to limit chronic remodeling processes may therefore translate into improved long-term function and reduced morbidity in children and adolescents.

More interestingly, bone marrow is classically considered an acute-responding tissue because of its high proliferative activity, with radiation-induced depletion typically manifesting rapidly and recovering over time. Yet, even after 10 weeks—when marrow repopulation would be expected—the FLASH group retained significantly higher marrow cellularity than the CONV group. This persistence suggests that FLASH not only attenuates the severity of initial depletion but also protects the regenerative capacity of the marrow niche, leading to durable preservation of hematopoietic function. Clinically, this finding is particularly important for pediatric osteosarcoma patients, who rely on intact marrow reserves to tolerate intensive, multi-agent chemotherapy. By sparing bone marrow while also protecting bone and muscle, FLASH may thus help maintain systemic treatment options and functional integrity, while allowing delivery of curative radiation doses.

The mechanisms underlying the FLASH effect remain incompletely understood but may involve transient oxygen depletion, altered redox signaling, modulation of immune responses, and protection of stem cell compartments. Our observation of preserved trabecular bone architecture and marrow cellularity suggests that FLASH may spare progenitor niches within the bone marrow microenvironment, thereby maintaining hematopoietic reserve. Reduced fibrosis and collagen deposition in muscle following FLASH parallels prior work linking diminished TGF-β1 signaling to mitigation of radiation-induced fibrosis in skin and other tissues (25). The protection observed in growing bones may also reflect enhanced remodeling capacity in juvenile animals, which could be preferentially preserved under FLASH conditions. Future studies incorporating molecular analyses, such as markers of osteoclast activity, osteoblast differentiation, and cytokine signaling, will be important to delineate the mechanistic basis of these observations.

Several strengths of this study should be noted. First, it represents the initial demonstration of the proton FLASH effect in juvenile animals, addressing a critical gap in modeling pediatric patients. Second, we provide long-term (10-week) data on bone, marrow, and muscle sparing, focusing on tissues most vulnerable in pediatric osteosarcoma treatment. Third, this is the report to validate synchrotron-based proton FLASH irradiation, demonstrating its feasibility and reproducibility. Because synchrotron accelerators remain integral to many hospital-based proton therapy centers worldwide, our findings significantly broaden the potential clinical applicability of FLASH.

This work also has limitations. We employed a single high dose (11 Gy) rather than a fractionated regimen, and additional studies are required to determine whether the FLASH effect is maintained under hypo fractionated or conventional fractionation schemes relevant to clinical practice. The 10-week follow-up period, while sufficient to capture late toxicities, does not encompass lifelong skeletal development, which is particularly relevant in pediatrics. Our endpoints were primarily structural and histological; future work should include assessments of fracture incidence, marrow lineage recovery, and molecular biomarkers of fibrosis and bone turnover. We did not include tumor-bearing models in this study and therefore cannot confirm that tumor control is preserved in juvenile osteosarcoma under FLASH conditions. Finally, the sample size, while sufficient to detect statistically significant differences, remains modest, and validation in larger cohorts will be necessary.

Despite these limitations, the clinical implications of our findings are substantial. For pediatric osteosarcoma, where curative radiotherapy is often constrained by the risk of late toxicity including growth impairment, fractures, and marrow suppression, proton FLASH offers the potential to deliver higher doses required for local control while sparing musculoskeletal development. This may expand the role of radiotherapy as a definitive modality for unresectable or anatomically complex tumors, reducing reliance on morbid surgery. Our results also provide proof-of-concept that FLASH platforms can achieve biologically meaningful tissue sparing, supporting their use in future clinical feasibility trials. Ongoing investigations should extend follow-up, incorporate tumor-bearing juvenile models, and evaluate the dependence of the FLASH effect on LET variations across the SOBP.

## Conclusion

In conclusion, proton FLASH irradiation confers significant long-term protection to bone, marrow, and muscle in juvenile mice, establishing both biological relevance and technical feasibility. By demonstrating durable musculoskeletal sparing in a pediatric-relevant model, this study provides critical preclinical evidence to support clinical translation of proton FLASH as a strategy to reduce morbidity and improve outcomes in pediatric osteosarcoma.

This study demonstrates that synchrotron-based proton FLASH irradiation significantly reduces chronic toxicities in bone, bone marrow, and muscle in juvenile mice compared with CONV irradiation. Using microCT and histological analyses, we observed preservation of bone mineral density, trabecular structure, marrow cellularity, and muscle integrity 10 weeks after exposure. These findings provide the first evidence that the FLASH effect can be reproduced in a synchrotron platform and, importantly, extend its protective benefits to the developing skeleton and musculoskeletal tissues most vulnerable in pediatric patients.

Our results complement prior work performed in adult animal models with cyclotron-based systems and uniquely establish the biological relevance of proton FLASH for pediatric osteosarcoma. By mitigating late musculoskeletal toxicities, synchrotron-based proton FLASH may enable escalation of therapeutic doses while preserving growth, marrow reserve, and long-term function. These findings strongly support continued investigation of FLASH in juvenile, tumor-bearing osteosarcoma models and motivate future studies exploring fractionation, LET dependence, and clinical feasibility.

Together, this work highlights proton FLASH as a compelling strategy to expand the therapeutic window of radiotherapy and improve outcomes for pediatric osteosarcoma patients with unresectable or high-risk diseases.

## Declaration of generative AI and AI-assisted technologies in the manuscript preparation process

During the preparation of this work the author(s) used ChatGPT in order to improve the language. After using this tool/service, the author(s) reviewed and edited the content as needed and take(s) full responsibility for the content of the published article.

## Conflict of Interest Statement for All Authors

None

## Funding Statement

This research is funded by Internal Seed Grant at MD Anderson Cancer Center

## Data Availability Statement for this Work

Research data are stored in an institutional repository and will be shared upon request to the corresponding author.

## Acknowledgements

We would like to thank the MD Anderson Small Animal Imaging Facility (SAIF), supported by the NIH/NCI under award number P30CA016672, for providing microCT scanning and image reconstruction. We thank the Hitachi team for their technical support in operating the synchrotron beamline and for providing reliable proton FLASH output for this study. We also acknowledge the DVMS Pathology Core at MD Anderson Cancer Center for their expert assistance with tissue processing, including paraffin embedding, sectioning, staining, and slide scanning.

## Notes

### Competing Interest Statement

The authors have declared no competing interest.

